# Genome-wide patterns of regulatory divergence revealed by introgression lines

**DOI:** 10.1101/037333

**Authors:** Rafael F. Guerrero, Amanda L. Posto, Leonie C. Moyle, Matthew W. Hahn

**Affiliations:** Department of Biology, Indiana University, Bloomington IN 47405, USA; School of Informatics and Computing, Indiana University, Bloomington IN 47405, USA

**Keywords:** Gene expression, *cis*-regulation, *trans*-regulation, hybrid misregulation, *Solanum lycopersicum*, *Solanum pennella*

## Abstract

Understanding the genetic basis for changes in transcriptional regulation is an important aspect of understanding phenotypic evolution. Using interspecific introgression lines, we infer the mechanisms of divergence in genome-wide patterns of gene expression between the nightshades *Solanum pennellii* and S. *lycopersicum* (domesticated tomato). We find that *cis-* and trans-regulatory changes have had qualitatively similar contributions to divergence in this clade, unlike results from other systems. Additionally, expression data from four tissues (shoot apex, ripe fruit, pollen, and seed) suggest that introgressed regions in these hybrid lines tend to be down-regulated, while background (non-introgressed) genes tend to be up-regulated. Finally, we find no evidence for an association between the magnitude of differential expression in NILs and previously determined sterility phenotypes. Our results contradict previous predictions of the predominant role of *cis-* over trans-regulatory divergence between species, and do not support a major role for gross genome-wide misregulation in reproductive isolation between these species.

## Introduction

Transcriptional regulation plays an important role in evolution (reviewed in Wray et al. 2003; Wittkopp 2013). Changes in the timing and level of gene expression, resulting from the evolution of cis-regulatory DNA sequences and trans-acting factors, underlie many phenotypic changes and can contribute to reproductive isolation between species (*e.g*., Jones *et al*. 2012; Scarpino *et al*. 2013; Turner *et al*. 2014). While the molecular basis for *cis*- and *trans*-acting changes is fairly well understood (*e.g*., Gibson and Weir 2005), the relative contribution of these mutations to the evolution of gene expression remains unknown. Because *cis*-regulatory regions are thought to be more robust to mutation (Ludwig *et al*. 2005) and less prone to deleterious pleiotropic effects (Prud’homme *et al*. 2007), cis-acting mutations have been predicted to accumulate at a faster rate than their *trans* counterparts (Wray 2007; Wittkopp *et al*. 2008). As a result of their faster fixation rate, cis-regulatory differences may compose a larger fraction of regulatory divergence. However, this prediction has received conflicting support: while some studies have found that most regulatory differences between species involve cis-acting mutations (Tirosh *et al*. 2009), others have found the opposite pattern (McManus *et al*. 2010; Meiklejohn *et al*. 2014). Moreover, in one case the contribution of cis-mutations did not seem to increase with divergence time (Coolon *et al*. 2014).

In order to infer the genetic basis of changes in gene expression, researchers have largely used two complementary approaches. One approach treats expression levels as quantitative traits, and then carries out standard QTL mapping in the context of a recombinant population (*e.g*., Schadt *et al*. 2003). These experiments are often referred to as “expression QTL” or “eQTL” studies. An advantage of this approach is that single loci can be identified that affect expression at multiple genes, thus helping to uncover the number of targeted genes for *trans*-acting factors, and possibly also the degree of pleiotropy of each. A disadvantage is that eQTLs that co-localize with their targets could be due to either cis-acting mutations or nearby trans-acting regulators (Rockman and Kruglyak 2006), and it is therefore difficult to estimate the exact contribution of *cis*-regulatory changes in these studies. The second approach to exploring regulatory divergence is to study gene expression patterns of inter-population or inter-specific hybrids. In *F*_1_ hybrids, inferences of *cis* and *trans* divergence can be made by contrasting allelic expression levels to those of the parental species (Wittkopp *et al*. 2004). This approach explicitly allows researchers to disentangle the effects of *cis*- and trans-acting changes on a single locus, though no inferences can be made concerning the number or identity of targets of trans-acting factors.

Interspecific hybrids, such as the ones generated by these experimental approaches, might also reveal the buildup of genetic incompatibilities in their genome-wide patterns of gene expression (Landry *et al*. 2007). Hybrids frequently show transgressive expression (*i.e*., outside the range of the parents), potentially as the result of incompatibilities in regulatory elements (*e.g*., faulty interactions between a transcription factor from one parent and a regulatory sequence from the other). Some studies have suggested, after finding increased levels of transgression in sterile hybrids, that pervasive misregulation contributes to hybrid sterility (Michalak and Noor 2003; Moehring *et al*. 2007; Ortíz-Barrientos *et al*. 2007; Rottscheidt and Harr 2007). However, while this association has been found in some systems (*Mus*, (Good *et al*. 2010; Turner *et al*. 2014); *Xenopus*, (Malone *et al*. 2007)), a relationship between misregulation and sterility is absent in others (*Drosophila*, (Barbash and Lorigan 2007); *Arabidopsis*, (Walia *et al*. 2009)). To generate a more general picture of regulatory divergence, and its potential role in hybrid sterility, data from a broader range of species and hybrid genotypes is needed.

An alternative to comparing expression patterns in parental genotypes to early-generation hybrids is to systematically assess gene expression in more targeted hybrid regions, using nearly isogenic lines (NILs; also known as introgression lines) between divergent genotypes (as in Meiklejohn *et al*. 2014). Such lines carry a short, often homozygous, chromosomal region from one genotype (species) on the background of a second genotype (species). Compared to *F*_1_ hybrids or *F*_2_ mapping populations, examining gene expression in NILs allows us to observe the regulatory machinery of the genotypic background acting on a limited number of genes from a second genotype (*i.e* those contained within the introgressed region), as well as the downstream effects of those introgressed loci on the background genotype (Fig. 1A). For genes that are differentially expressed between parental genotypes, introgressed loci can be examined for whether expression levels resemble either parent, are intermediate (not significantly different from either), or are transgressive. In introgressed genes, expression resembling the donor parent suggests a stronger effect of cis-acting (or tightly linked) elements. In contrast, expression levels similar to those of the recipient parent indicate the action of trans-regulatory factors, while intermediate or transgressive expression levels imply the interaction of both mechanisms. When NILs are available for the majority of a genome, there is the opportunity to broadly examine the prevalence of each class of interactions genome-wide. Therefore, NILs represent an opportunity to gain a comprehensive view of the genome-wide prevalence of *cis*-versus *trans*-regulation, as well as potential regulatory incompatibilities when crosses are made between species.

**Figure 1.**
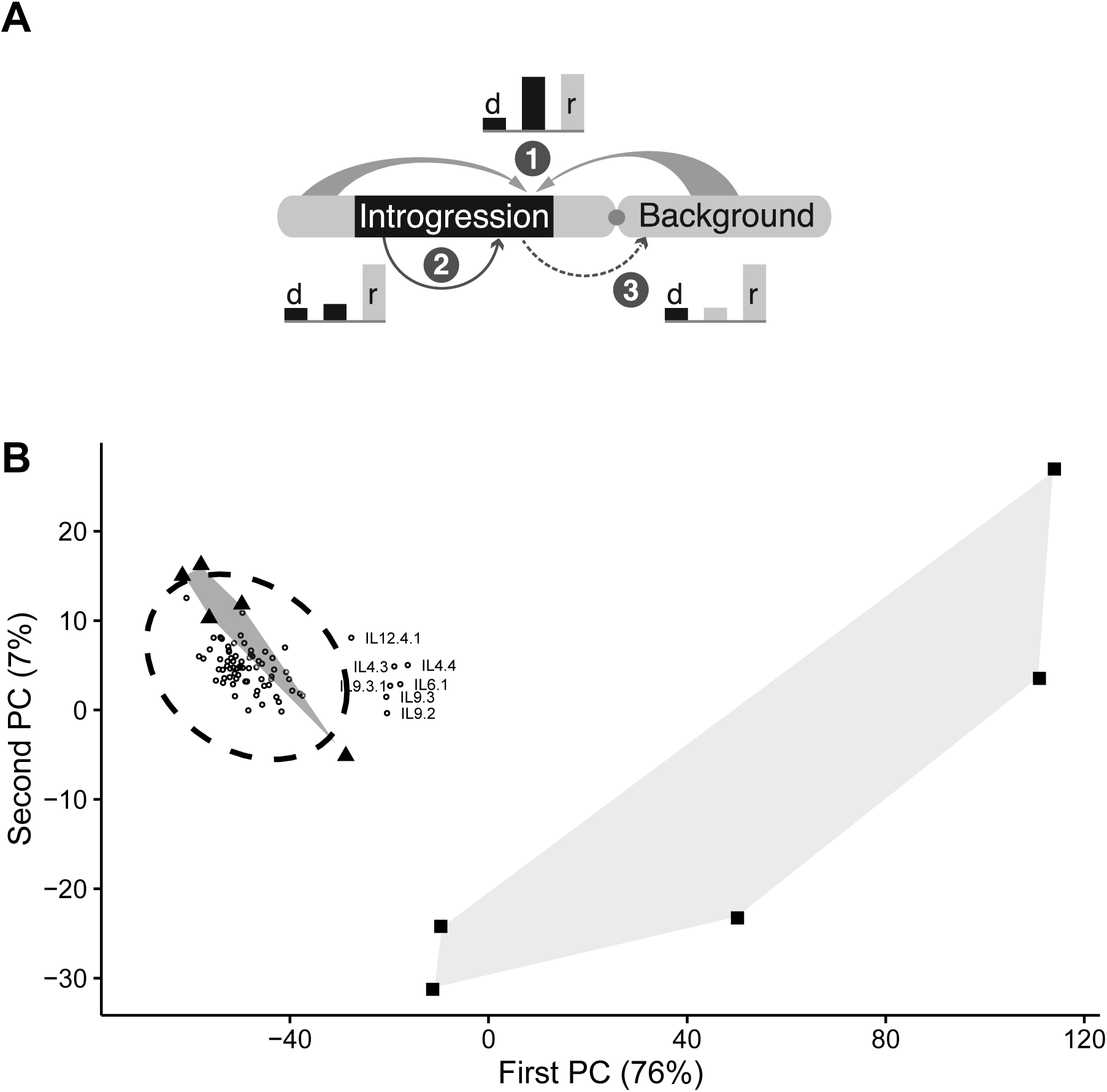
A) Expression of introgressed genes in NILs is the result of the interaction between (1) *trans*-regulatory factors from the recipient parent and (2) putatively cis-regulatory elements in the introgressed region. The introgression, in turn, may affect background expression (3). For genes with divergent expression between parents (bars, labeled ‘d’ for donor and ‘r’ for recipient), we expect expression in introgressions to be recipient-like (1, middle bar) if divergence has been predominantly *trans*. On the other hand, cis-regulatory divergence is expected to result in donor-like expression in introgressed regions. B) Genome-wide regulatory divergence between the recipient (*S. lycopersicum*; triangles) and the donor parent (*S. pennellii*; squares) summarized in the first two eigenvectors of a principal components analysis. Expression profiles of NILs (open circles) projected on to these principal components show that introgression lines have expression patterns closer to the recipient parent. Only the mean for each NIL is shown, and the dashed ellipse represents the 95% confidence interval of the distribution of all NILs (assuming a multivariate t-distribution). The shaded areas representing the expression range of the parents are drawn for clarity only.

The domesticated tomato (*Solanum lycopersicum)* and its wild relative, *S. pennellii,* present an excellent pair of species in which to study the evolution of regulatory divergence. These two South American nightshades are close relatives (about 1.5% sequence divergence; Pease *et al*. 2016), but show considerable differences in their gene expression profiles (Koenig *et al*. 2013). Although the two species are differentiated by loci that cause partial pollen and seed sterility ((Nakazato *et al*. 2008), and see below), they can produce hybrids, which has allowed for the development of nearly 80 NILs. Each of these hybrid lines carries, in homozygous form, an introgression of *S. pennellii* on a *S. lycopersicum* background (Eshed and Zamir 1994, 1995). The average introgressed region comprises about 700 genes (~2% of the *S. lycopersicum* genome), and 97% of known genes have been introgressed in at least one NIL.

Genome-wide expression levels of these NILs have not been examined in the context of understanding the genome-wide prevalence of *cis*- and *trans*-regulation. Here, we use gene expression profiles of NILs to study the mechanisms underlying regulatory evolution. We infer the relative roles of *cis*- and *trans*-regulatory divergence, and find that comparable fractions of genes are predominantly influenced by each mechanism. Additionally, because we are interested in the potential relationship between high levels of misregulation and hybrid sterility, we characterize gene expression levels of seven NILs that have previously shown to have sterility phenotypes (Moyle and Nakazato 2008). In contrast to some previous studies, we find no evidence of increased magnitude of misregulation in sterile tissues, suggesting that sterility in this system is not related to detectable regulatory incompatibilities.

## Methods

To describe the patterns of gene expression in tomato NILs and the relative contributions of *cis* and *trans* divergence, we analyzed publicly available RNA-seq libraries from vegetative tissue (Chitwood *et al*. 2013). To explore the correlation between levels of misregulation and sterility, we carried out a microarray hybridization experiment on sterile pollen and seed tissues. Additionally, we used publicly available data from fruit tissue to confirm some of our inferences. The analysis of this third, more limited, dataset is reported in the Supplement.

To describe the genome-wide regulatory effects of introgression, we counted genes with significant differences in expression level with respect to the recipient parent, *S. lycopersicum*. We refer to this count as the differential expression level, and to these genes as “differentially expressed.” We use the amount of differential expression in the background as a proxy for hybrid misregulation (*sensu* Barbash and Lorigan 2007). We quantified levels of differential expression in introgressed and background regions of each NIL. To determine which genes reside in introgressed regions, we used the described genomic boundaries of introgressions (from Chitwood *et al*. 2013) and gene models in the ITAG 2.3 tomato genome (available from the SOL Genomics Network, solgenomics.net; Mueller *et al*. 2005). We assessed whether introgressed regions had significantly more differentially expressed genes by evaluating whether they were overrepresented in the genome-wide counts of differentially expressed genes per NIL, using Fisher’s exact tests.

We used the vegetative tissue gene expression levels to infer underlying regulatory mechanisms. We classify NIL gene expression as “donor-like” (*i.e*., resembling donor parent *S. pennellii*), “recipient-like” (*i.e*., resembling recipient parent *S. lycopersicum*), “intermediate,” or “transgressive” based on three pairwise contrasts between donor, recipient, and NIL. For “divergent” genes (*i.e*., those with significant differences between the parental lines), expression in NILs is considered donor-like if it differed significantly from the recipient but not the donor. If the opposite is true, expression is classified as recipient-like. Genes are classified as intermediate if their expression is not significantly different from either parent, or if it is significantly different from both parents but occurs within the parental range. A gene is classified as transgressive if its expression level is outside the parental range and is differentially expressed with respect to both parents.

To compare downstream effects of introgressions across NILs we used a negative binomial linear model *D_b_* ∽ *D_i_* + *N* + *S* + *∊*, where *D_b_* is the differential expression level in the background, *D_i_* is the differential expression level in the introgressed region, *N* is the total number of introgressed genes, and *S* is a sterility state. Lines were categorized as “sterile” if they carry sterility QTL (18 NILs; Moyle and Nakazato 2008), or “non-sterile” otherwise (see Supplemental Table 8). This allowed us to evaluate the relationship between misregulation and sterility while controlling for NIL size.

All analyses were done in R (R Core Team 2015), and the scripts used are included in the Supplement.

### Expression in vegetative tissue

We analyzed gene expression in shoot apices (including young leaves) from RNA-seq libraries published in Chitwood *et al*. (2013). In that study, the two parental lines and 76 NILs were exposed to two experimental conditions. Here we limit our analyses to the libraries from the “sun” treatment, as this is the more standard growth condition (we also ran the analysis using the “shade” subset and found similar patterns; results not shown). We analyzed five biological replicates per genotype (*i.e*., parent or NIL), for a total of 390 libraries. We use the counts of reads obtained by Chitwood *et al*. (2013), who mapped each NIL to a custom reference genome based on the *S. lycopersicum* genome (ITAG version 2.3; 34727 gene models) but included the sequence of *S. pennellii* at the corresponding introgressed region. This approach minimizes the possibility of mapping bias due to sequence divergence. In particular, this approach allows us to exclude the possibility of apparent down-regulation of *S. pennellii* alleles as a result of transcript sequence differences between species (which could lead to a failure to map *S. pennellii* reads onto a *S. lycopersicum* reference genome). We filtered out genes with low expression levels, keeping those that show more than two read-counts per million in at least five libraries (using the *edgeR* package; (Robinson *et al*. 2010)). A total of 20110 genes were analyzed in this tissue, of which 19460 are introgressed at least once (96.8%). Genes may be introgressed in multiple NILs: in this dataset, 7755 genes are introgressed in only one of the 76 NILs, 9233 are introgressed in two different NILs, 2121 genes in three NILs, and 351 in four NILs.

We carried out a *voom* transformation of the raw counts (Law *et al*. 2014) and fit a linear model using the *lmFit* function (Smyth 2004) from the *limma* package (Ritchie *et al*. 2015). We used the linear model *ϒ* ~ *A* + ∊, where *ϒ* is the normalized expression level, and *A* is the accession of the sample (*A* has 78 levels: 76 NILs and two parents). We computed t-statistics with empirical Bayes adjustment of standard errors using the *eBayes* function (Law *et al*. 2014). In order to get the significance of the pairwise comparisons between parents and each NIL, we used the *decideTests* function from the same package to get the significance of the contrasts of interest, controlling for a false discovery rate of 5%.

### Expression in sterile tissues

We selected three NILs with reduced seed set (IL1.1, IL1.4, and IL2.3) and four with reduced pollen fertility (IL1.1.3, IL4.2, IL7.2, and IL8.1.1), based on known QTL for these sterility phenotypes (Moyle and Nakazato 2008). Eight individuals per NIL and twelve individuals of *S. lycopersicum* were grown in a randomized common garden experiment to flowering. We quantified gene expression using an Affymetrix tomato GeneChip®. We analyzed a total of 38 microarray hybridizations: six biological replicates for each tissue in *S. lycopersicum*, and four replicates per NIL [with the exception of IL1.1.3 (pollen) and IL1.4 (seed) for which we used three replicates per NIL].

#### Plant material

All plants were propagated under the same conditions, following standard greenhouse cultivation protocols (Moyle and Graham 2005). Seeds were germinated and transplanted as seedlings into flats. Three weeks after transplant, seedlings were transferred to individual 3.75L pots and grown in a climate-controlled greenhouse at the Indiana University Biology Greenhouse facility. Plants were watered daily, fertilized weekly, and staked prior to flowering. Collected tissue was frozen in liquid nitrogen and stored at -80°C until RNA was extracted. Anthers were collected from the third flower on an inflorescence at preanthesis (*i.e*., the day before the flower opened), when normally developing pollen is known to be mature but not yet dehisced. Carpels were collected seven days after pollination. To do so, the third flower on an inflorescence was emasculated prior to anthesis and immediately pollinated with pollen freshly collected from another flower on the same individual. Hand-pollinated flowers were tagged prior to collection.

#### RNA isolation

We used a Trizol (Invitrogen) extraction followed by RNeasy purification (Qiagen). Anthers and carpels from each sample were individually homogenized in liquid nitrogen and dissolved in Trizol at a ratio of 1 mL/100 mg Trizol to tissue. After the addition of chloroform (1/5 TRIzol volume) and centrifugation for phase separation, the aqueous phase containing RNA was removed, the volume was recorded, and 0.53X volumes of 100% ethanol was added. The mixture was then applied to an RNeasy mini column (QIAGEN) and purified RNA was obtained following the manufacturer’s instructions. For the seed sterility experiment, 2 carpels were pooled per biological replicate. For the pollen sterility experiment, 1-3 anthers were pooled per biological replicate. For some biological replicates, tissue was pooled from multiple individuals to obtain sufficient RNA yield (no individuals were used in more than one pool). Samples were pooled following individual extractions and RNA was quantified prior to pooling using a Nanodrop ND-1000 as well as preparing a gel. We used 6 ug (for seed samples) and 7.5 ug (for pollen samples) of RNA for sample preparation and hybridizations.

#### Microarray sample preparation and hybridization

The Affymetrix tomato GeneChip was designed based on tomato Unigene Build #20, and GenBank mRNAs up to November 5, 2004. Sample preparation and hybridization was performed at the CMG (Center for Medical Genomics), IUPUI (Indianapolis). For each sample, single cycle labeling was performed followed by product purification, quantitation, and fragmentation. After adding control oligonucleotides, 200 uL of hybridization cocktail was applied to an array and incubated. Following incubation, washing and staining steps were performed in an Affymetrix Fluidics Station. To reduce non-random error, balanced groups of samples were handled in parallel. Arrays were scanned using a dedicated scanner, controlled by Affymetrix GCOS software, and images examined for defects. Microarray intensities (CEL files) were processed using the Affy package (Gautier *et al*. 2004) in R (R Core Team 2015).

#### Differential gene expression analysis

We include a total of 5228 singly-mapping probesets that target an equal number of genes. Mapping annotation of probesets to ITAG2.3 genes was downloaded from the SOL Genomics Network (Mueller *et al*. 2005). As in the vegetative tissue analysis, we carried out a *voom* transformation and fit a model using the *limma* package. For each tissue, we fit a linear model *ϒ* ~ *A* + ∊, where *ϒ* is the normalized expression level and *A* is the accession variable. In the pollen model, *A* has 5 levels (4 NILs and the recipient parent), while in the seed model it has 4 levels (3 NILs and the recipient parent). To account for multiple testing, we control the false discovery (FDR) at a 5% level in each tissue.

## Results

### Expression in vegetative tissue

We found that introgressed regions are over-represented for differentially expressed genes. Each NIL has on average 135 differentially expressed genes (ranging from 62 genes in lines IL4.1.1 and IL7.1, to 470 genes in IL4.4). Relative to their size, introgressed regions carry a large fraction of all differentially expressed genes in each NIL: about 30% of differentially expressed genes in NILs are located in introgressions, even though each individual introgression makes up on average 2% of the tomato genome. In fact, all NILs show a significant overrepresentation of differentially expressed genes in introgressed regions (Fisher’s exact test, *p* < 0.01 in all cases; Supplemental Table 8). In turn, this implies a relatively smaller effect of introgressed genes on the expression of background genes. On average, we estimate that each introgressed gene has downstream effects on the expression of about 0.4 genes outside the introgressed region. As a result, genome-wide expression profiles of NILs fall roughly within the range of the recipient parent (Fig. 1B). Seven NILs show profiles slightly more similar to the donor parent, falling outside of the 95% confidence interval of the estimated distribution across all NILs (labelled in Fig. 1B).

In contrast, the parents show sub-stantial differences in their expression profiles: we find 4251 divergent genes (FDR=5%), representing 21% of genes expressed in vegetative tissue. The observed divergence in gene expression has been reported previously (Koenig *et al*. 2013), and implies that low levels of differential expression in the NILs are unlikely to be due to lack of differences in the parents. We also found that the donor parent, the wild species *S. pennellii*, displays much more variation than the recipient parent, domesticated *S. lycopersicum*. This difference is consistent with higher levels of genome-wide heterozygosity in *S. pennellii* (Pease *et al*. 2016) and larger phenotypic variance in *S. pennellii* (Muir *et al*. 2014).

We found that differential expression of genes in background and introgressed regions differs not only in magnitude, but also in direction (Fig. 2). While introgressed regions show a trend towards down-regulation, differentially expressed genes in background regions of NILs tend to be up-regulated. As mentioned in the Methods, the mapping approach implemented ensures that down-regulation in introgressed regions is unlikely to be an artifact of interspecific sequence mismatch. Moreover, most differentially expressed genes in introgressed regions vary in the direction of the donor (*i.e*., are donor-like or intermediate), while very few are transgressive. In contrast, a larger fraction of differentially expressed genes in the background is transgressive. This large fraction of genes with expression levels outside of the parental range is likely the result of the joint effects of the compensatory mutations underlying *cis*-by*-trans* interactions (as in Landry *et al*. 2005) and/or the indirect effects of the introgression (*e.g*., the down-regulation of an introgressed gene leads to failed repression of another gene in the network).

**Figure 2.**
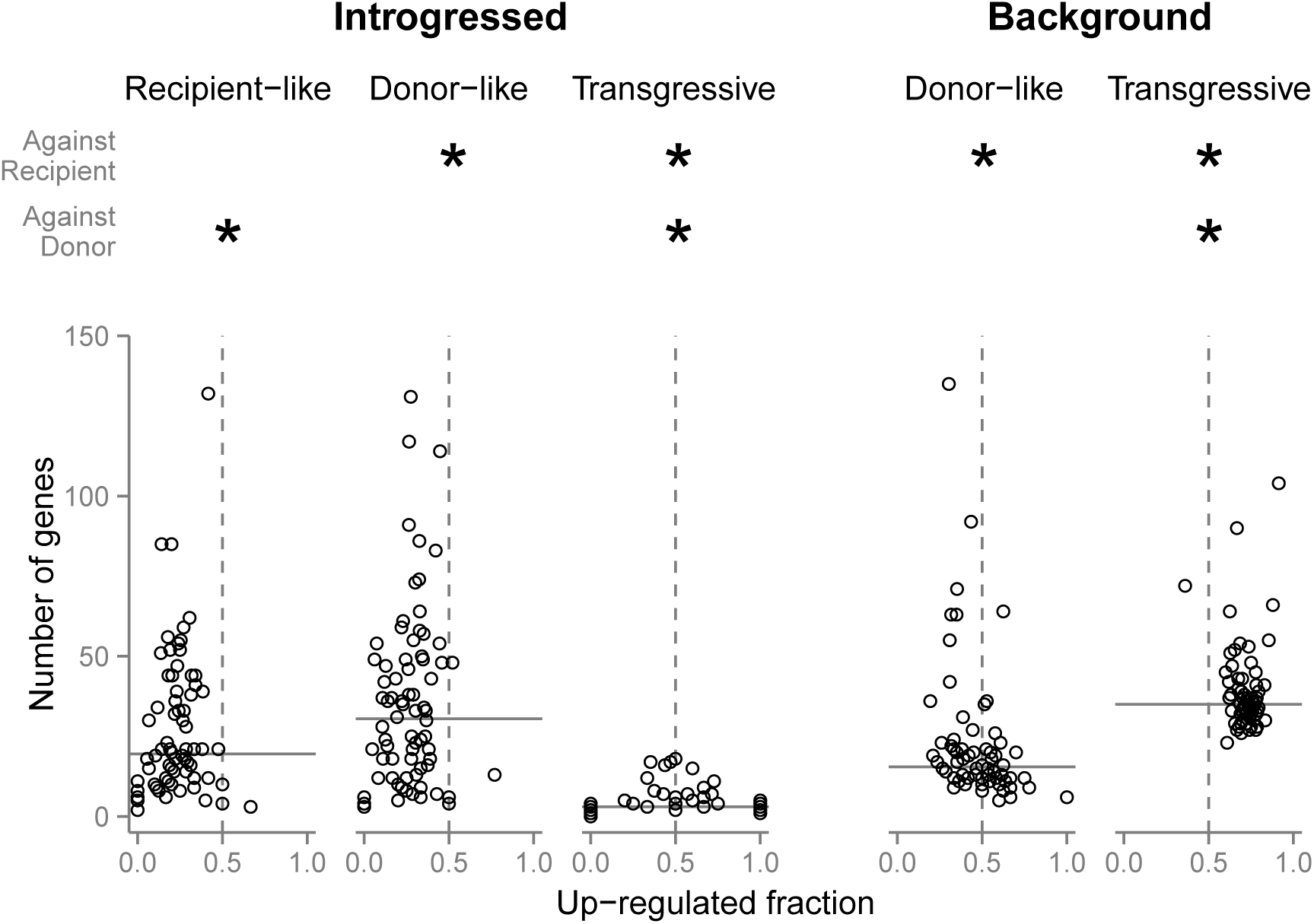
The number of differentially expressed genes by NIL, and the fraction of those genes that are up-regulated, varies between regions of the genome. Transgressive genes compose a large fraction of the differentially expressed genes in background regions, but not in introgressed regions. Down-regulated genes are more common in introgressed donor-like genes, but more transgressive genes in the background show up-regulation

### Relative contributions of *cis* and *trans* divergence

Overall, expression patterns suggest that *cis*- and trans-regulatory changes have contributed equally to divergence between *S. lycopersicum* (the recipient parent) and *S. pennellii* (the donor parent) (Fig. 3). A total of 4277 introgressed genes show differences in expression that allow us to make inferences about the predominant regulatory mechanisms. First we restrict our inferences to 3366 genes (79% of those differentially expressed) that have consistent expression patterns across NILs—*i.e.*, those that can be classified in the same category every time they appear in introgressions (Fig. 3). We find that a considerable fraction (36%) of divergent genes show donor-like expression levels. Comparable fractions of introgressed genes show intermediate (30%) or recipient-like expression (28%). Additionally, we find evidence of transgressive expression for 6% of examined genes. Of the 911 introgressed genes that show inconsistent expression patterns across NILs, 79% vary between resembling either parent (*i.e*., donor- or recipient-like expression) and showing intermediate levels, while 18% vary between donor-like and recipient-like expression (Supplemental Table 2).

**Figure 3.**
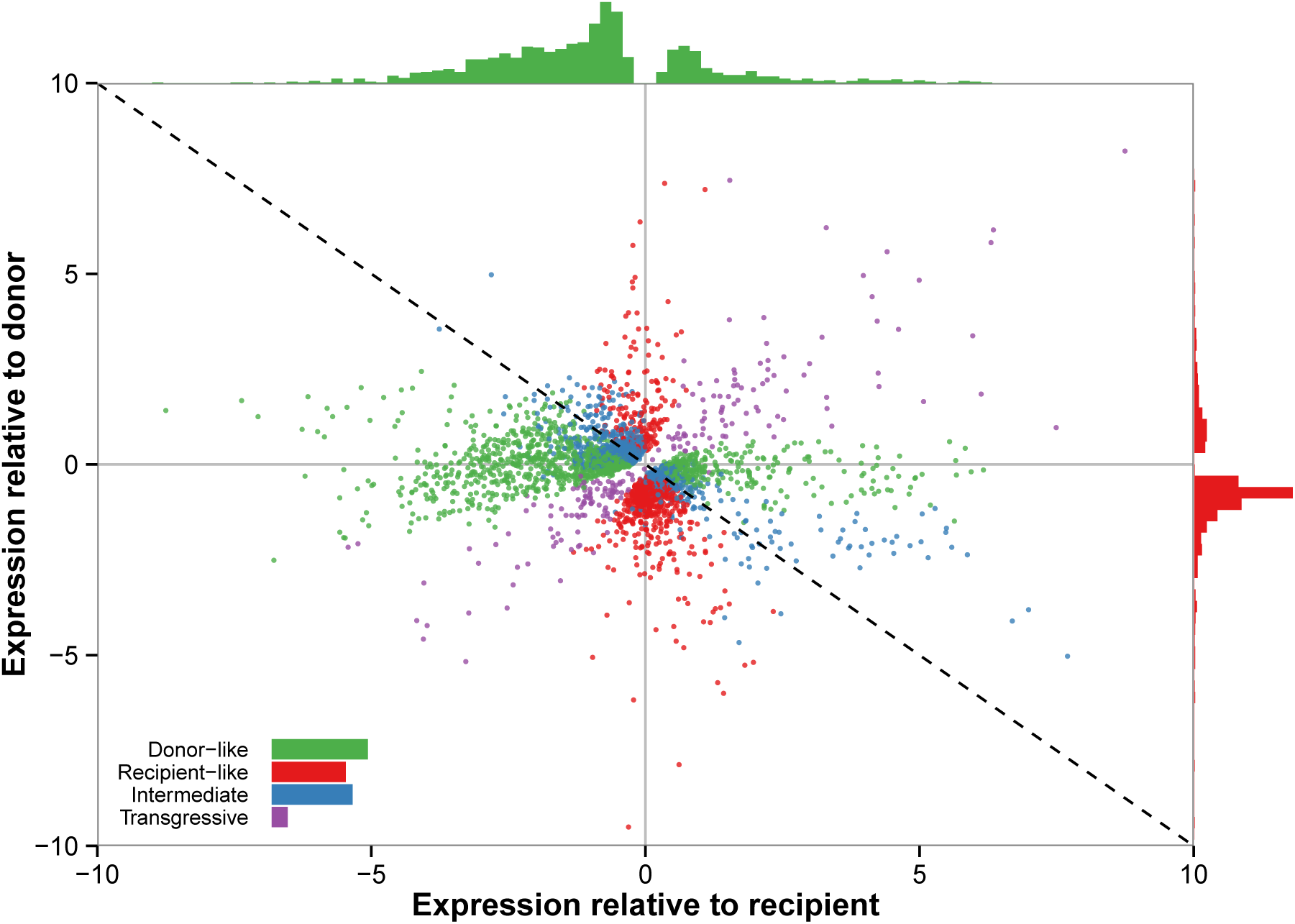
Normalized expression levels for 3366 introgressed genes expressed in vegetative tissue, relative to both parents. Many introgressed genes show donor-like expression (36%; green points), but comparable fractions of genes are observed for intermediate (30%; blue points) and recipient-like expression (28%; red points). The distribution of expression levels relative to the recipient parent for donor-like genes (green histogram, top) shows a bias towards down-regulation. On the right (red histogram), a similar pattern for expression levels relative to the donor parent in recipient-like genes.

These results imply that there is an approximately equal contribution of *cis*- and *trans*-regulatory mutations to the observed divergence in expression levels. In introgressed regions, *cis*-regulatory donor alleles interact with the recipient *trans*-regulatory network from the background genome. Therefore, donor-like expression of genes inside the introgression suggests that the causal mutation for expression divergence is linked to the introgressed gene. Conversely, recipient-like expression of introgressed alleles indicates a more prominent role of *trans*-acting elements in the regulatory divergence of those genes. The large fraction of genes with intermediate or transgressive levels suggests that linked (putatively cis-acting) and unlinked (*trans*-acting) elements contribute significantly to the resulting expression profile. Interestingly, we find a trend towards down-regulation of introgressed alleles, both in donor-like and recipient-like genes (Fig. 3), suggesting that interaction of regulatory elements in hybrids is more likely to result in failure to express rather than overexpression, regardless of the nature of the causal mutation.

### Expression in sterile tissues

Differential expression patterns in sterile pollen and seeds are similar to those found in vegetative tissue (and fruits, see Supplement). Specifically, we observe an excess of differentially expressed genes in introgressed regions (Fisher’s exact, *p* < 0.01 in all cases). Again, we see a trend towards down-regulation in introgressed regions and up-regulation in background genes (Fig. 4). These qualitative similarities suggest that increased differential expression is not a distinctive feature of sterile hybrid genotypes, and therefore imply that gross misregulation patterns in hybrids do not play a major role in reproductive incompatibility in these species. The observed over-representation of differential expression in introgressions is unlikely to be the product of differential hybridization to the microarray due to sequence divergence, since we find that the result is still significant if we consider only up-regulated genes (Fisher’s exact, *p* < 0.05 in all but one NIL; Supplemental Table 7). Moreover, we observe similar results if we limit the analysis to probes with identical sequences in the two parents (not shown).

**Figure 4.**
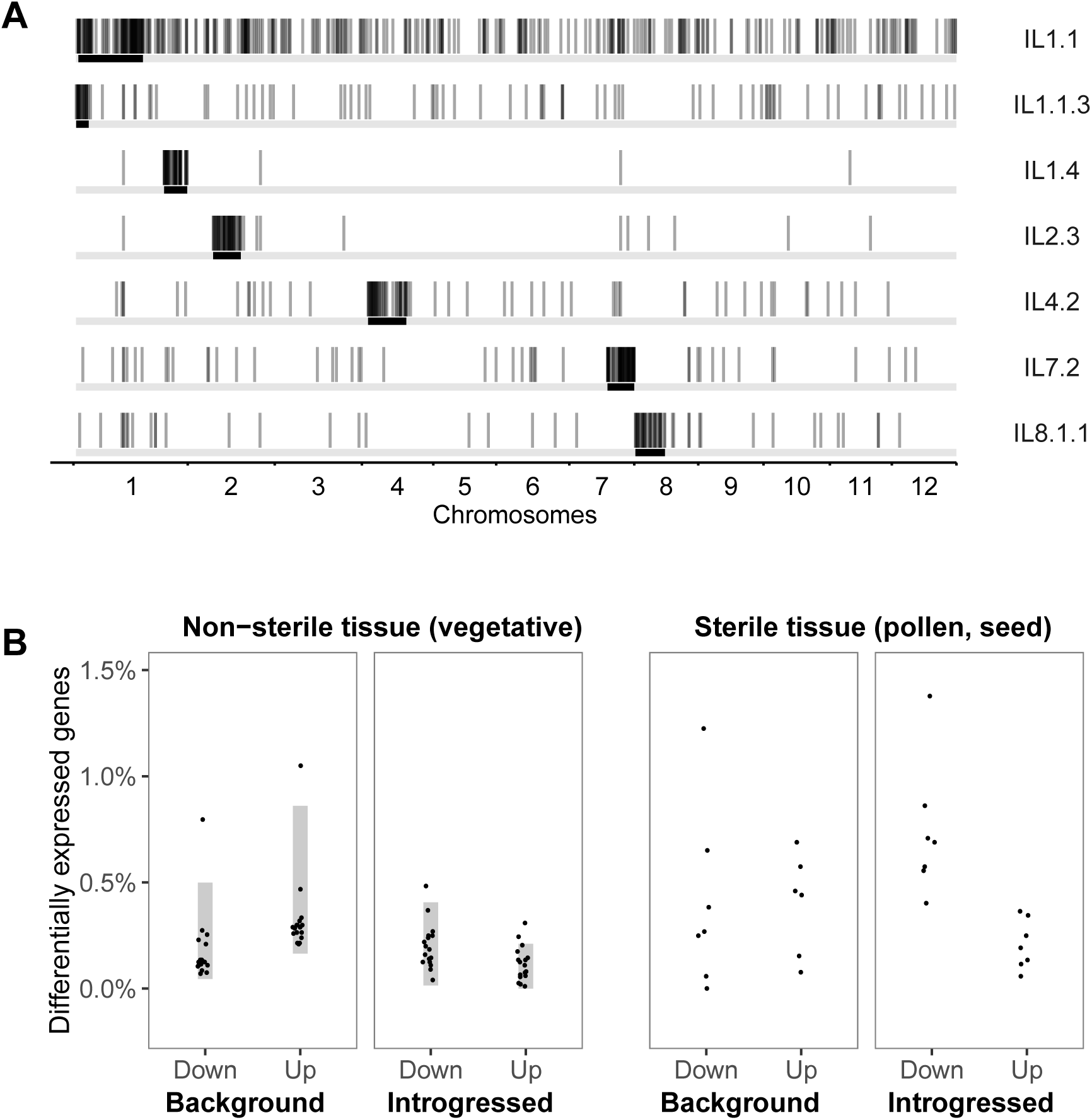
A) Genomic location of differentially expressed genes (vertical bars) in sterile tissue of seven NILs. Most differential expression accumulates in introgressed regions (black area in horizontal lines). B) No qualitative difference in the differential expression patterns of sterile compared to non-sterile tissue of NILs that carry sterility QTL, as described by the percent of total number of genes queried in each tissue (5228 genes in sterile tissue, 20110 in vegetative tissue). The gray vertical bars represent the 95% CI for 61 NILS that do not carry known sterility QTL.

Consistent—albeit indirectly—with this result, NILs that carry sterility QTLs do not show increased differential expression in vegetative tissue (Fig. 4). Despite sub-stantial variation between NILs in the amount of background differential expression (ranging from 49 genes in introgression line IL1.2 to 397 genes IL4.4), there is no evidence that differential expression is correlated with sterility phenotypes. In particular, the differential expression levels in eighteen NILs associated with sterility phenotypes are not significantly greater than in other NILs (in vegetative tissue or fruits; Supplemental Tables 1, 5). One NIL linked to seed infertility (IL4.4), however, does show the highest number of differentially expressed background genes in vegetative tissue (but it is not an outlier in fruit; see Supplement). Although we originally included IL4.4 in our microarray experiment, we were unable to generate sufficient post-fertilization material so we cannot determine whether the sterility QTL in it has a large effect on seed gene expression.

## Discussion

The expression profiles of NILs can be used to reveal the mechanisms behind the evolution of gene regulation in their parental species. In *Solanum*, the most common pattern was for genes to show donor-like expression, suggesting an important role of cis-regulatory divergence in this system. However, sub-stantial fractions of genes also show evidence of *trans-* (recipient-like, 28%) or *cis-by-trans* regulatory divergence (intermediate or transgressive, together 36%). Moreover, not all genes that show donor-like expression are necessarily cis-regulated, as some of them could be targets of trans-regulatory elements residing in the same NIL (Rockman and Kruglyak 2006). These linked (but trans-acting) elements may inflate our estimate of the contribution of *cis*-regulatory divergence. Given these results, we conclude that *cis* and *trans* mechanisms participate approximately equally in this species comparison.

Our results are consistent with previous findings in *Coffea* (Combes *et al*. 2015) and *Arabidopsis* (Shi *et al*. 2012), where divergence between species is only slightly biased towards cis-acting mutations. By contrast, most studies in *Drosophila* suggest a stronger role of cis-regulation, but there is considerable variation in the results (reviewed in Coolon and Wittkopp 2013). In fact, divergence between *D. mauritiana* and *D. simulans* is mostly trans-regulatory (Meiklejohn *et al*. 2014). Interestingly, in that system Meiklejohn *et al*. 2014 found that the effect of introgression on background differential expression (which they interpret as trans-regulation) is biased towards up-regulation. Our results are consistent with their findings, and together they allow us to speculate that a large fraction of trans-regulatory divergence involves disrupted interactions between negative regulators and their targets.

Our results (as well as others; Shi *et al*. 2012; Meiklejohn *et al*. 2014; Combes *et al*. 2015), disagree with the proposed prevalence of *cis*- over trans-regulatory divergence between species (Wray *et al*. 2003; Wray 2007; Wittkopp *et al*. 2008). The basis of this prediction is intuitive: mutations in *trans*-acting elements should be, on average, more deleterious than those in *cis*-regulatory sequences because they are more likely to have negative pleiotropic effects. Therefore, natural selection should fix a higher proportion of *cis*- relative to *trans*-regulatory mutations. Over time, the different evolutionary dynamics of the two types of mutations lead to a proportional increase of *cis*-acting mutations contributing to regulatory divergence. Thus, an underlying assumption made when studying regulatory differences between species is that enough time has passed to allow for detectable over-accumulation of *cis* divergence, but this need not be the case. Moreover, the proportions of *cis*- and *trans-* variants segregating in the ancestral population are unknown. Elevated *trans* variation in the ancestor (as in Emerson *et al*. 2010; Goncalves *et al*. 2012) will increase *trans* divergence, and more time would be necessary to fix enough *cis*- regulatory mutations to observe a bias towards *cis* divergence.

The expectation of increase in *cis* divergence through time may not hold due to at least three other factors. First, *cis*- and *trans*-acting mutations may be equally constrained if *cis*-acting mutations have *trans*-acting consequences downstream in the gene network, resulting in similar pleiotropic constraints on both mutation types. Second, the prediction assumes equal opportunity for *cis* and *trans* mutations; however, gene regulation is commonly polygenic, and in species with complex regulatory networks the increased opportunity for *trans*-regulatory mutations (as in (Gruber *et al*. 2012)) may outweigh the negative skew in the distribution of their fitness effects. Greater opportunities for trans-acting mutations could mean that we observe more of them, even if on average they are more deleterious than cis-acting mutations. Third, the deleterious pleiotropic effects of *trans* mutations may be somewhat reduced in systems with redundant or context-dependent regulation. Further study of the molecular basis behind trans-regulation in this system (e.g., the role of chromatin modification, small RNAs) will be necessary to understand the causes underlying the genome-wide regulatory patterns we observe.

We found no relationship between the magnitude of misregulation and hybrid sterility, consistent with previous findings in *Drosophila* (Barbash and Lorigan 2007). In contrast, mouse trans-eQTLs of large effect co-localize with sterility QTL, suggesting that quantitative misregulation plays a role in hybrid incompatibility in that system (Turner *et al*. 2014). In tomatoes, introgressed regions act as trans-acting factors for the differentially expressed genes in the background (*i.e*., they are trans-eQTLs), and some of these introgressions carry sterility QTL (Moyle and Nakazato 2008). However, expression levels in sterile tissues of these NILs do not show dramatic differences with respect to those of non-sterile tissues. Moreover, we find no co-localization of trans-eQTL and sterility loci, as sterile NILs show similar levels of misregulation as other lines in vegetative tissue (Fig. 4). This result does not imply that regulatory incompatibilities do not play a role in reproductive isolation between these two species. Rather, our results indicate that the high number of misregulated genes *per se* is not the cause of low fitness in hybrids.

Here, we contribute to an expanding body of literature that, taken together, does not support a difference in the evolutionary dynamics of *cis*- and *trans*-regulatory mechanisms. While multiple factors—such as demography (McManus *et al*. 2010), natural history of the species (Combes *et al*. 2015), or methodological issues (Meiklejohn *et al*. 2014)—may be invoked to explain why previous studies have not found the predicted excess of *cis* over *trans* divergence, we believe that we now have enough evidence to question the generality of such expectations.

## Acknowledgments

We thank Steven Dunkelbarger, Sarah Josway, and Emily Josephs for help with plant cultivation and materials, and Simo Zhang and James Pease for technical help during data analysis. We also thank the reviewers for their helpful comments. We are grateful for funding support from NSF DEB-0532097, DEB-0841957, and MCB-1127059, as well as the Indiana Metabolomics and Cytomics Initiative (METACyt).

## References

Barbash, D. A., and J. G. Lorigan. 2007. Lethality in *Drosophila melanogaster/Drosophila simulans* species hybrids is not associated with sub-stantial transcriptional misregulation. Journal of Experimental Zoology Part B: Molecular and Developmental Evolution 308B:74–84.

Chitwood, D. H., R. Kumar, L. R. Headland, A. Ranjan, M. F. Covington, Y. Ichihashi, D. Fulop, J. M. Jiménez-Gómez, J. Peng, J. N. Maloof, and N. R. Sinha. 2013. A quantitative genetic basis for leaf morphology in a set of precisely defined tomato introgression lines. The Plant Cell 25:2465–2481.

Combes, M.-C., Y. Hueber, A. Dereeper, S. Rialle, J.-C. Herrera, and P. Lashermes. 2015. Regulatory divergence between parental alleles determines gene expression patterns in hybrids. Genome Biology and Evolution 7:1110–1121.

Coolon, J. D., C. J. McManus, K. R. Stevenson, B. R. Graveley, and P. J. Wittkopp. 2014. Tempo and mode of regulatory evolution in *Drosophila*. Genome res 24:797–808.

Coolon, J. D., and P. J. Wittkopp. 2013. *cis*- and trans-Regulation in *Drosophila* Interspecific Hybrids. Pp. 37–57 in. Chen, Z. J., and J. A. Birchler, eds. Polyploid and Hybrid Genomics. John Wiley & Sons, Inc.

Emerson, J. J., L. Hsieh, H. Sung, T. Wang, C. Huang, H. H. Lu, M. J. Lu, S. Wu, and W. Li. 2010. Natural selection on *cis* and *trans* regulation in yeasts. Genome res 20:826–836.

Eshed, Y., and D. Zamir. 1994. A genomic library of *Lycopersicon pennellii* in *L. esculentum:* a tool for fine mapping of genes. Euphytica 79:175–179.

Eshed, Y., and D. Zamir. 1995. An introgression line population of *Lycopersicon pennellii* in the cultivated tomato enables the identification and fine mapping of yield-associated QTL. Genetics 141:1147–1162.

Gautier, L., L. Cope, B. M. Bolstad, and R. A. Irizarry. 2004. affy—analysis of Affymetrix GeneChip data at the probe level. Bioinformatics 20:307–315.

Gibson, G., and B. Weir. 2005. The quantitative genetics of transcription. Trends Genet 21:616–23.

Goncalves, A., S. Leigh-Brown, D. Thybert, K. Stefflova, E. Turro, P. Flicek, A. Brazma, D. T. Odom, and J. C. Marioni. 2012. Extensive compensatory *cis-trans* regulation in the evolution of mouse gene expression. Genome res 22:2376–2384.

Good, J. M., T. Giger, M. D. Dean, and M. W. Nachman. 2010. Widespread overexpression of the X chromosome in sterile F1 hybrid mice. PLos Genet 6:e1001148.

Gruber, J. D., K. Vogel, G. Kalay, and P. J. Wittkopp. 2012. Contrasting properties of gene-specific regulatory, coding, and copy number mutations in *Saccharomyces cerevisiae*: Frequency, effects, and dominance. PLos Genet 8:e1002497.

Jones, F. C., M. G. Grabherr, Y. F. Chan, P. Russell, E. Mauceli, J. Johnson, R. Swofford, M. Pirun, M. C. Zody, S. White, et al. 2012. The genomic basis of adaptive evolution in threespine sticklebacks. Nature 484:55–61.

Koenig, D., J. M. Jiménez-Gómez, S. Kimura, D. Fulop, D. H. Chitwood, L. R. Headland, R. Kumar, M. F. Covington, U. K. Devisetty, and A. V. Tat. 2013. Comparative transcriptomics reveals patterns of selection in domesticated and wild tomato. Proc Natl Acad Sci U S A 110:2655–2662.

Landry, C., D. Hartl, and J. Ranz. 2007. Genome clashes in hybrids: insights from gene expression. Heredity 99:483–493.

Landry, C. R., P. J. Wittkopp, C. H. Taubes, J. M. Ranz, A. G. Clark, and D. L. Hartl. 2005. Compensatory cis-trans evolution and the dysregulation of gene expression in interspecific hybrids of *Drosophila*. Genetics 171:1813–1822.

Law, C. W., Y. Chen, W. Shi, and G. K. Smyth. 2014. Voom: precision weights unlock linear model analysis tools for RNA-seq read counts. Genome Biol 15:R29.

Ludwig, M. Z., A. Palsson, E. Alekseeva, C. M. Bergman, J. Nathan, and M. Kreitman. 2005. Functional evolution of a cis-regulatory module. PLoS Biol 3:e93.

Malone, J. H., T. H. Chrzanowski, and P. Michalak. 2007. Sterility and gene expression in hybrid males of *Xenopus laevis* and *X. muelleri*. PLoS One 2:e781.

McManus, C. J., J. D. Coolon, M. O. Duff, J. Eipper-Mains, B. R. Graveley, and P. J. Wittkopp. 2010. Regulatory divergence in *Drosophila* revealed by mRNA-seq. Genome Res 20:816–825.

Meiklejohn, C. D., J. D. Coolon, D. L. Hartl, and P. J. Wittkopp. 2014. The roles of *cis*- and trans-regulation in the evolution of regulatory incompatibilities and sexually dimorphic gene expression. Genome Res 24:84–95.

Michalak, P., and M. A. Noor. 2003. Genome-wide patterns of expression in *Drosophila* pure species and hybrid males. Mol Biol Evol 20:1070–1076.

Moehring, A. J., K. C. Teeter, and M. A. Noor. 2007. Genome-wide patterns of expression in *Drosophila* pure species and hybrid males. II. Examination of multiple-species hybridizations, platforms, and life cycle stages. Mol Biol Evol 24:137–145.

Moyle, L. C., and E. B. Graham. 2005. Genetics of hybrid incompatibility between *Lycopersicon esculentum* and *L. hirsutum*. Genetics 169:355–373.

Moyle, L. C., and T. Nakazato. 2008. Comparative genetics of hybrid incompatibility: sterility in two *Solanum* species crosses. Genetics 179:1437–1453.

Mueller, L. A., T. H. Solow, N. Taylor, B. Skwarecki, R. Buels, J. Binns, C. Lin, M. H. Wright, R. Ahrens, and Y. Wang. 2005. The SOL Genomics Network. A comparative resource for Solanaceae biology and beyond. Plant Physiol 138:1310–1317.

Muir, C. D., J. B. Pease, and L. C. Moyle. 2014. Quantitative Genetic Analysis Indicates Natural Selection on Leaf Phenotypes Across Wild Tomato Species (Solanum sect. Lycopersicon; Solanaceae). Genetics 198:1629–1643.

Nakazato, T., M. Bogonovich, and L. C. Moyle. 2008. Environmental factors predict adaptive phenotypic differentiation within and between two wild Andean tomatoes. Evolution 62:774–792.

Ortíz-Barrientos, D., B. A. Counterman, and M. A. Noor. 2007. Gene expression divergence and the origin of hybrid dysfunctions. Genetica 129:71–81.

Pease, J. B., D. C. Haak, M. W. Hahn, and L. C. Moyle. 2016. Phylogenomics reveals three sources of adaptive variation during a rapid radiation. PLoS Biology in press.

Prud’homme, B., N. Gompel, and S. B. Carroll. 2007. Emerging principles of regulatory evolution. Proc Natl Acad Sci U S A 104:8605–8612.

R Core Team. 2015. R: A Language and Environment for Statistical Computing. URL http://www.R-project.org/.

Ritchie, M. E., B. Phipson, D. Wu, Y. Hu, C. W. Law, W. Shi, and G. K. Smyth. 2015. limma powers differential expression analyses for RNA-sequencing and microarray studies. Nucleic Acids Res:gkv007.

Robinson, M. D., D. J. McCarthy, and G. K. Smyth. 2010. edgeR: a Bioconductor package for differential expression analysis of digital gene expression data. Bioinformatics 26:139–140.

Rockman, M. V., and L. Kruglyak. 2006. Genetics of global gene expression. Nat Rev Genet 7:862–872.

Rottscheidt, R., and B. Harr. 2007. Extensive additivity of gene expression differentiates subspecies of the house mouse. Genetics 177:1553–1567.

Scarpino, S. V., P. J. Hunt, F. J. Garcia-De-Leon, T. E. Juenger, M. Schartl, and M. Kirkpatrick. 2013. Evolution of a genetic incompatibility in the genus *Xiphophorus*. Mol Biol Evol 30:2302–2310.

Schadt, E. E., S. A. Monks, T. A. Drake, A. J. Lusis, N. Che, V. Colinayo, T. G. Ruff, S. B. Milligan, J. R. Lamb, G. Cavet, P. S. Linsley, M. Mao, R. B. Stoughton, and S. H. Friend. 2003. Genetics of gene expression surveyed in maize, mouse and man. Nature 422:297–302.

Shi, X., D. W. Ng, C. Zhang, L. Comai, W. Ye, and Z. J. Chen. 2012. *cis*- and *trans*-regulatory divergence between progenitor species determines gene-expression novelty in *Arabidopsis* allopolyploids. Nat Commun 3:950.

Smyth, S. 2004. Linear models and empirical bayes methods for assessing differential expression in microarray experiments. Stat. Appl. Genet. Mol. Biol 3.

Tirosh, I., S. Reikhav, A. A. Levy, and N. Barkai. 2009. A yeast hybrid provides insight into the evolution of gene expression regulation. Science 324:659–662.

Turner, L. M., M. A. White, D. Tautz, and B. A. Payseur. 2014. Genomic networks of hybrid sterility. PLos Genet 10:e1004162.

Walia, H., C. Josefsson, B. Dilkes, R. Kirkbride, J. Harada, and L. Comai. 2009. Dosage-dependent deregulation of an AGAMOUS-LIKE gene cluster contributes to interspecific incompatibility. Curr Biol 19:1128–1132.

Wittkopp, P. J. 2013. Evolution of gene expression. Pp. 413–419 in. Losos, J. B., D. A. Baum, D. J. Futuyma, H. E. Hoekstra, R. E. Lenski, A. J. Moore, C. L. Peichel, D. Schluter, and M. C. Whitlock, eds. The Princeton guide to evolution.

Wittkopp, P. J., B. K. Haerum, and A. G. Clark. 2004. Evolutionary changes in *cis* and *trans* gene regulation. Nature 430:85–88.

Wittkopp, P. J., B. K. Haerum, and A. G. Clark. 2008. Regulatory changes underlying expression differences within and between *Drosophila* species. Nat Genet 40:346–350.

Wray, G. A. 2007. The evolutionary significance of cis-regulatory mutations. Nat Rev Genet 8:206–216.

Wray, G. A., M. W. Hahn, E. Abouheif, J. P. Balhoff, M. Pizer, M. V. Rockman, and L. A. Romano. 2003. The evolution of transcriptional regulation in eukaryotes. Mol Biol Evol 20:1377–1419.

